# Identification of dysregulation of atrial proteins in rats with chronic obstructive apnea using twodimensional polyacrylamide gel electrophoresis and mass spectrometry

**DOI:** 10.1101/388751

**Authors:** Jacob C. Lux, Devika Channaveerappa, Roshanak Aslebagh, Timothy A. Heintz, Meredith McLerie, Brian K. Panama, Costel C. Darie

**Affiliations:** Biochemistry and Proteomics Group, Department of Chemistry and Biomolecular Science, Clarkson University, Potsdam, NY 13699-5810; Department of Experimental Cardiology, Masonic Medical Research Laboratory, Utica, New York, USA 13501

**Keywords:** animal proteomics, rat, cardiovascular system, hypoxia, metabolism, two-dimensional electrophoresis, apnea

## Abstract

Obstructive sleep apnea (OSA) affects an estimated 20% of adults worldwide with up to 80% of patients remaining undiagnosed. OSA has been associated with electrical and structural abnormalities of the atria, although the molecular mechanisms are not well understood. We have implemented a rat model of OSA involving the surgical implantation of a tracheal obstructive device. Rats were divided into severe and moderate apnea groups, receiving 23 seconds (severe) or 13 seconds (moderate) apneas per minute, 60 apneas per minute for 8 hours a day over 2 weeks. We recently performed a pilot study using onedimensional polyacrylamide gel electrophoresis (1D PAGE) and nanoliquid chromatography-tandem mass spectrometry (NanoLC-MS/MS) to investigate the protein dysregulations in rat atria which was induced with OSA using the rat model we developed. We found, among others, that some aerobic and anaerobic glycolytic enzymes and Krebs cycle enzymes were downregulated, suggesting that apnea may be a result of paucity of oxygen and production of ATP and reducing equivalents. Here, we used twodimensional polyacrylamide gel electrophoresis (2D PAGE) coupled with nanoLC-MS/MS as a complementary approach to investigate the proteins that are dysregulated in the atria from severe and moderate apnea when compared to control. We not only found that the entire glycolytic pathway and Krebs cycle are downregulated, but also found evidence that additional enzymes involved in the beta-oxidation, electron transport chain and Krebs cycle anaplerotic reactions were also downregulated. Other protein dysregulations identified are involved in metabolic, structural, or inflammatory pathways, suggesting that these proteins may play a role in atrial pathology developing via chronic obstructive apnea and hypoxia.

## Introduction

Obstructive sleep apnea (OSA) is characterized by repeated cessation in respiration (apnea) in the upper airway during sleep lasting at least 10 seconds. Apneas occur due to temporary pharyngeal collapse or narrowing, resulting in decreased blood oxygen saturation [1]. OSA severity is measured via the apnea-hypopnea index (AHI), a calculation of the number of apneas (complete collapse of the airway) or hypopneas (partial collapse of the airway) the sufferer has per hour [2]. Mild OSA (AHI ≥5) affects about 1 in 5 adults and severe OSA (AHI ≥15) affects 1 in 15 adults, with an estimated 80% of sufferers being undiagnosed [2, 3]. OSA has been associated with headache, daytime sleepiness, obesity, depression, arthritis, type 2 diabetes mellitus, arteriosclerosis, atherosclerosis, hypertension, atrial arrhythmia, and sudden cardiac death [1, 2].

Although OSA has been closely associated with atrial arrhythmia, atrial enlargement, increased P wave duration, and decreased atrial voltage; little is known about the molecular pathways causing these and other pathologies [4–6]. Many animal models do not address both the obstructive and hypoxemic components of OSA in conscious, free roaming animals. Some current OSA models employ chronic intermittent hypoxia. While this causes drastic hypoxic swings, changes in oxyhemoglobin levels are not as rapid as in OSA patients. These models also do not reproduce the intrathoracic pressure swings seen in patients with OSA [7].

In order to accurately reproduce the effects of OSA as observed in the clinic, we have employed a surgical model involving a silicone obstructive device implanted in the trachea of conscious, free roaming rats [5, 8]. We have found this model to accurately replicate rapid oxygen desaturation seen in the clinic while applying complete tracheal occlusion. In our previous 1D-PAGE study, we found significant dysregulations for metabolic proteins, suggesting a decrease in glycolysis and a diminished ability to produce reducing equivalents for ATP generation. Structural sarcomere proteins were also up-regulated, which is consistent with cardiac hypertrophy. Our results pointed towards the changes that occur with OSA and how they contribute to the development of cardiac disease.

Here, we used 2D PAGE coupled with nanoLC-MS/MS as a complementary approach to investigate the proteins that are dysregulated in the atria from severe and moderate apnea when compared to control. While the 1D-PAGE approach compared the whole atrial proteomes from the severe OSA, moderate OSA and controls, the 2D-PAGE approach allowed us to identify only the dysregulated proteins from these conditions. Furthermore, 2D-PAGE also allowed us to identify different protein isoforms, already demonstrated in others’ work [9, 10].

In the 2D-PAGE experiments, we not only found that the entire glycolytic pathway and Krebs cycle are downregulated, but also found evidence that additional enzymes involved in the beta-oxidation, electron transport chain and Krebs cycle anaplerotic reactions were also downregulated. Other protein dysregulations identified are involved in metabolic, structural, or inflammatory pathways, suggesting that these proteins may play a role in atrial pathology developing via chronic obstructive apnea and hypoxia.

## Materials and Methods

### OSA Model

The use of 50-70 day old Sprague Dawley rats conformed to the Guide for the Care and Use of Laboratory Animals, published by the National Institutes of Health [11]. All research protocols were approved by the Animal Care and Use Committee at the Masonic Medical Research Laboratory. The rat model setup was as described previously [5]. Briefly, rats were anesthetized using isoflurane (5% for induction, 1.5-2% for maintenance). A 2.5 cm midline incision was made in the ventral portion of the neck and blunt dissection used to expose the trachea. The obstruction tubing was twined through the holes and secured in place with two 6-0 sutures. A second incision was made between the scapulae and the PE50 tubing was tunneled subdermally to the posterior incision. The sternohyoid muscle and dermis were closed with sutures and triple antibiotic applied topically (Actavis, Parsippany, NJ, USA). Rats were administered buprenorphine (30 μg/kg) for pain management immediately after surgery. Clavamox (40 mg/kg) was administered on a 5 day regimen, beginning the day before surgery.

Rats were allowed one week of recovery while attached to a tether (SAH-18; SAI) and PE50 tubing was attached to the L-shaped tube between the scapulae. Rats were free-roaming and had access to food and water during the tethered rest period and the two week apnea administration. Apneas lasting 13 or 23 seconds were delivered to rats in the moderate and severe apnea groups, respectively. Apneas (inflations) were delivered randomly via computer-controlled compressed air system (**Fig. 1**). The rats were given 13 or 23 second apneas 60 times per hour for 8 hours in a day. Apnea induction was confirmed using both MouseOx pulse oximetry (Starr Life Sciences, Oakmont, PA, USA) and direct observation of increased respiratory effort upon device inflation. Electrocardiogram (ECG) readings were also recorded. Control rats received tracheal surgery, but no device inflations.

**Fig 1.**
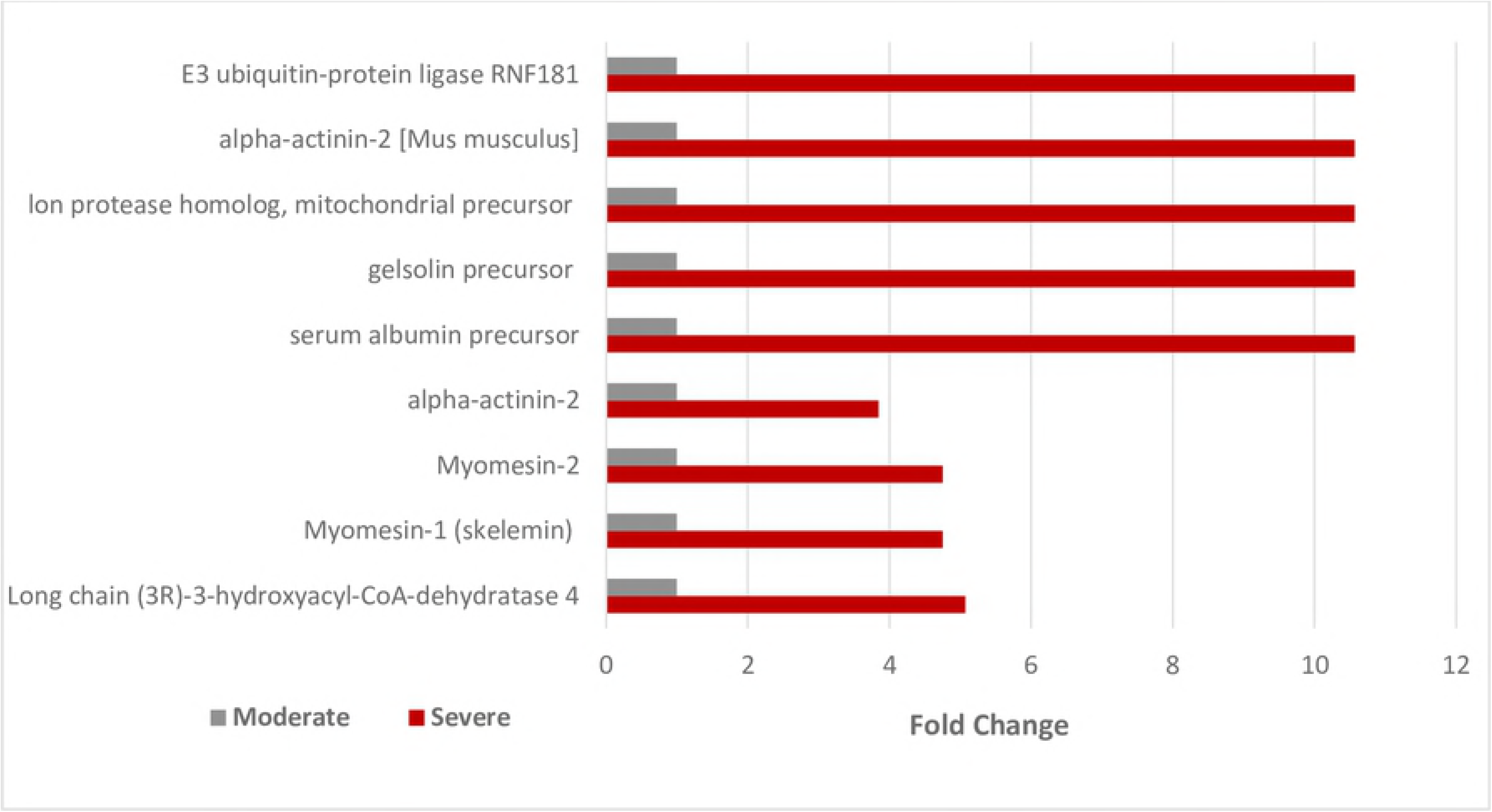
Illustration of OSA setup. Rats were tethered to a computer controlled compressed air system and allowed access to food and water.

### Sample preparation

Rats were anesthetized via IP injection of combination of ketamine (80 mg/kg) and xylazine (8 mg/kg). The rat hearts were rapidly removed, washed in chilled Tyrode’s solution, and was perfused according to the Langendorff technique. Atrial tissue was then flash frozen with liquid N_2_ and ground with a mortar and pestle. Tissue was placed in 300 μl of lysis solution containing NaCl 100 mM, Tris-HCl 25 mM, EDTA 0.2 mM, NaF 2 mM, Na_3_VO_4_ 2 mM and Complete Protease Inhibitor (Sigma, St. Louis, MO, USA). The lysed tissue was homogenized by passing the solution through sterile 18 and 22 gauge needles. Homogenate was centrifuged at 16,000 g for 5 min. and the supernatant collected and placed on ice. The pellet was resolubilized in 300 μl of lysis buffer with 2% NP40. Solution was incubated on ice for one hour and centrifuged at 16,000 g for 5 min. The supernatant was set aside on ice and the pellet was once again resolubilized in 200 μl of lysis buffer with 4% SDS and centrifuged at 16,000 g for 5 min. The supernatants were combined and protein quantification was done using bicinchoninic acid (BCA) assay (ThermoFisher Scientific, Waltham, MA, USA), and samples were stored at −80°C until use.

### Two Dimensional PAGE (2D-PAGE)

Atrial homogenates of control (n=2), moderate (n=2), and severe (n=2) apnea were used for analysis. For each sample condition and its biological replicate, coomassie and silver stained gels were obtained which were loaded with 550μg and 150μg of protein respectively. 2-D electrophoresis was performed by Kendrick Labs, Inc. (Madison, WI) according to the carrier ampholine method of Isoelectric focusing (IEF) [12–14]. IEF was performed in a glass tube with an inner diameter of 3.3 mm, using 2.0% pH 3-10 Isodalt Servalytes (Serva, Heidelberg, Germany) for 20,000 volt-hours. 100 ng of tropomyosin was added as IEF international standard to each sample. A pH electrode was used to determine the tube gel pH gradient plot for this set of Servalytes.

After 10 minutes of equilibration in 10% glycerol, 50mM dithiothreitol, 2.3% SDS and 0.0625 M tris, pH 6.8; each tube gel was sealed to the top of a stacking gel overlaying a 10% acrylamide slab gel (1.0 mm thick). SDS gel electrophoresis was performed over ~5 hours at 25 mA/gel. Myosin (220,000), phosphorylase A (94,000), catalase (60,000), actin (43,000), carbonic anhydrase (29,000), and lysozyme (14,000) were used as molecular weight standards and appear as bands on the basic edge of the silver stained 10% acrylamide gels. Gels were dried between cellophane paper sheets with the acid edge to the left.

### Computerized Comparisons

Gels were performed in duplicate and scanned with a laser densitometer (Model PDSI, Molecular Dynamics Inc, Sunnyvale, CA). The scanner was checked for linearity prior to scan with a calibrated Neutral Density Filter Set (Melles Griot, Irvine, CA). Progenesis Same Spots software version 4.5, 2011, TotalLab, UK) and Progenesis PG240 software (version 2006, TotalLab, UK) were used to analyze the images. Computerized analysis involved warping images, spot finding, background subtraction (average on boundary), matching, and quantification.

Spot % is equal to spot integrated density above background (volume). This was expressed as a percentage of total density above the background of all spots measured. The difference between corresponding spots from different sample condition (e.g. severe versus control) is expressed as fold-change of spot percentages. A positive fold change is indicative of the upregulation of the protein in the spot and a negative fold change indicates the downregulation of the protein in the spot.

### Spot Picking and In-Gel Digestion

A total of 2293 protein spots were compared for statistical differences, of which 208 spots had a fold increase or decrease of ≥1.7 and a P-value ≤0.05, or a fold increase or decrease of ≥3.0 were selected and excised for in-gel trypsin digestion and peptide extraction for nanoLC-MS/MS analysis. Excised spots were washed in HPLC grade water, 50 mM ammonium bicarbonate (ABC), 50% acetonitrile (ACN)/ 50% ABC for 30 minutes each under moderate shaking at room temperature. Gel pieces were dehydrated with 100% ACN and dried under speed vac. Dithiothreitol (DTT) (10 mM) and 25 mM ABC were used for reduction for 1 hour at 57° C. Iodoacetamide (100 mM) in 25 mM ABC was used for alkylation over 45 minutes, in the dark. Gel pieces were dehydrated again with 100% ACN and speed vac and then rehydrated with 20 μg of trypsin solution (10 ng/μl). Samples were incubated at 37° C overnight. After incubation, peptide extraction was performed with 5% formic acid (FA)/50 mM ABC/ 50% ACN and with 5% FA/100% ACN for 1 hour each. Peptides were dried and cleaned with a C18 Zip-Tip (reversed phase chromatographic column) (Millipore, Billerica, MA) and solubilized in 0.1% FA/2% ACN.

### NanoLC-MS/MS analysis and Data processing

The peptides were analyzed using a NanoAcquity UPLC (Waters Corp., Milford, MA, USA) coupled to a Q-TOF Xevo G2 MS (Waters, Milford, MA, USA). A C18 1.7 μm, 150 μm x 100 mm reversed phase column (Waters, Milford, MA, USA) was used to chromatographically separate the peptides over a 60 minute gradient of 1-85% ACN in 0.1% FA at a flow rate of 400 nl/min. MS/MS spectra were obtained in a data-dependent acquisition mode consisting of a survey of the five most intense peaks and automatic data-dependent MS/MS of 2+, 3+ 4+, 5+ and 6+ ions. MS/MS was triggered when MS signal intensity raised to 350 counts/seconds. Full MS scan covered the m/z range from 350 to 2000. Calibration was performed for precursor and product ions using 1 pmol GluFib standard peptide (Glu1-Fibrinopeptide B) with the amino acid sequence EGVNDNEEGFFSAR and a calculated mass for the monoisotopic m/z peak of 1570.68. The monitored precursor ion had an m/z of 785.84 (2+).

Raw data was processed using ProteinLynx Global Server (PLGS, version 2.4, Waters Corporation, Milford, MA, USA) software with the following parameters: background subtraction of polynomial order 5 adaptive with a threshold of 35%, two smoothings with a window of three channels in Savitzky–Golay mode, and centroid calculation of the top 80% of peaks based on a minimum peak width of 4 channels at half-height. The Mascot Daemon software (Matrix Science) was employed for database searches. Search parameters used were: propionamide as fixed modification for cysteine, methionine oxidation as variable modification, and precursor and product/ fragment mass tolerance were set to 0.5 Da and 0.8 Da respectively against enzyme trypsin with one missed cleavage. The pkl files with the mentioned parameters were search against NCBInr rat (*rattus*) database.

### Bioinformatics Analysis

The PANTHERv.13.1 (Protein Analysis Through evolutionary Relationships) classification system was used for the classification of proteins identified based on their molecular function of proteins interacting at a biochemical level and also on the biological processes [15, 16]

## Results and Discussion

Comparison of the protein pattern between severe versus control and moderate versus control was done using 2 biological replicates for each condition. Two Coomassie and 1 silver stained gels were run for each of the 6 samples (2 controls, 2 moderates and 2 severe samples). **Fig 2** shows an example of one Coomassie stained gel for each sample and **S1 Fig** shows the silver stained gel images. The two silver stained gels for each condition were averaged and the spot% was calculated. A total of 2293 spots were detected across averaged biological replicate gels of control, moderate, and severe 2D PAGE gels. **S2 Fig** depicts the 2D gel difference image of averaged severe apnea vs averaged control where the spots increased in severe are shown in blue and the spots decreased in severe are shown in red. Overall, 208 spots were selected for comparison if a p ≤0.05 and a fold change of ≥1.7 or if fold change was ≥3.0 were detected. 208 spots were in-gel digested using trypsin and were analyzed via Nano LC-MS/MS. Proteins were identified using the MASCOT Daemon database search engine. A summary and comparison of all the proteins identified from the spots of severe, moderate and control gels are shown in **S1-S3 Tables**. All the proteins listed in the three tables are only selected if the protein score (sum of the highest ions score for each distinct sequence) on MASCOT database was higher than 25 with a minimum of 2 peptide identifications for the protein.

**Fig 2.**
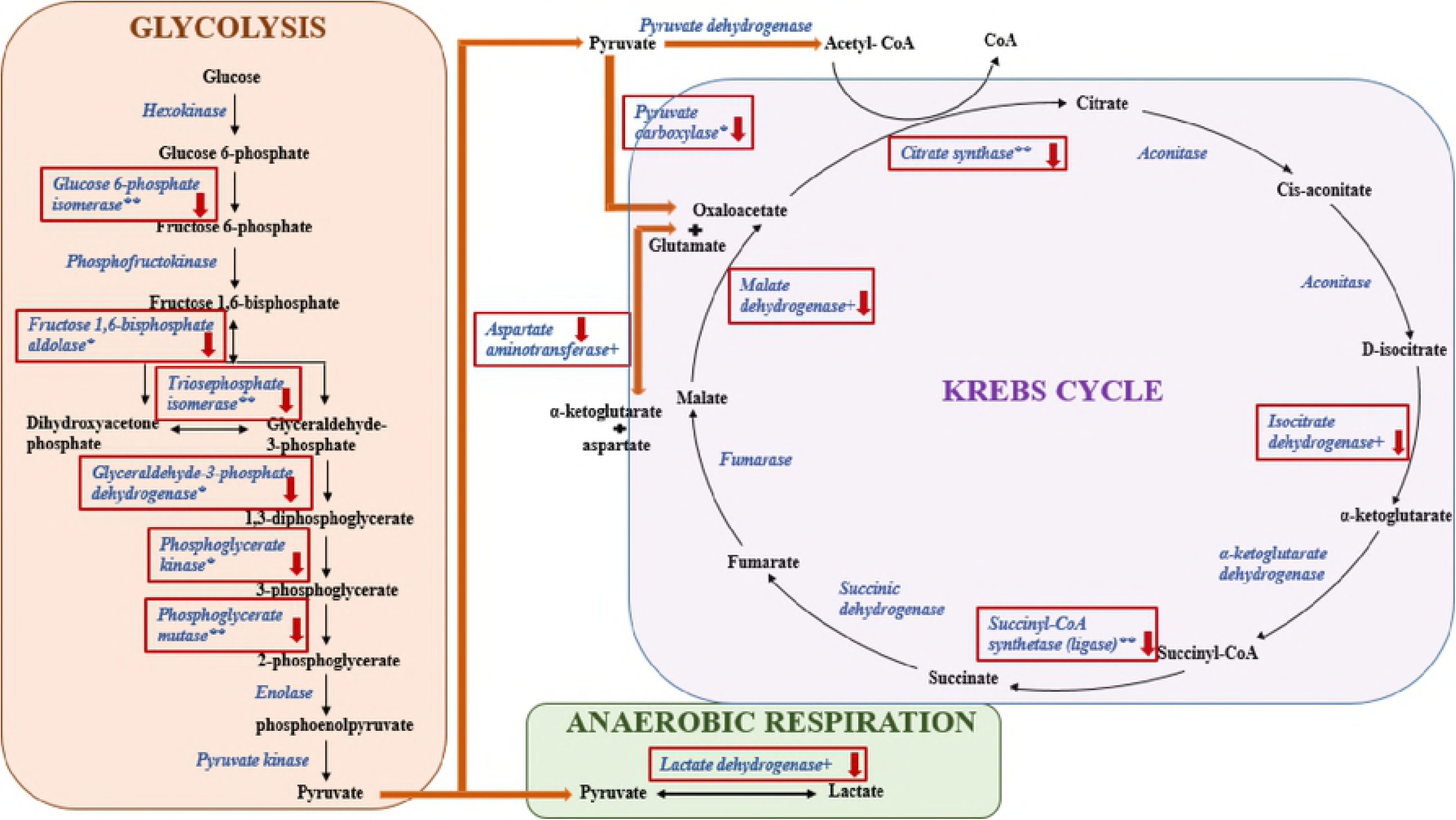
Images of control, moderate and severe apnea coomassie stained 2D polyacrylamide gels. The red circle on the left side of each polyacrylamide gel indicates the location of the IEF internal standard, tropomyosin with a Mr of 33,000 and pI of 5.2

### Dysregulated proteins in severe apnea samples, compared to controls

Upregulated proteins in severe apnea samples when compared to controls are shown in **Fig 3**. Some examples of the dysregulated proteins include creatine kinase M-type (9.8 fold), aldose reductase (9.8 fold), long-chain specific acyl-CoA dehydrogenase, mitochondrial precursor (9.8 fold), alpha-actinin-2 (6.7 fold), alpha-myosin heavy chain, partial (6.2 fold), myomesin-1 (skelemin) (4.8 fold), myosin light chain 4 (4.7 fold), myomesin-2 (4.6 fold); myosin light chain (4.0 fold) and heat shock protein 70 (3.9 fold).

**Fig 3.**
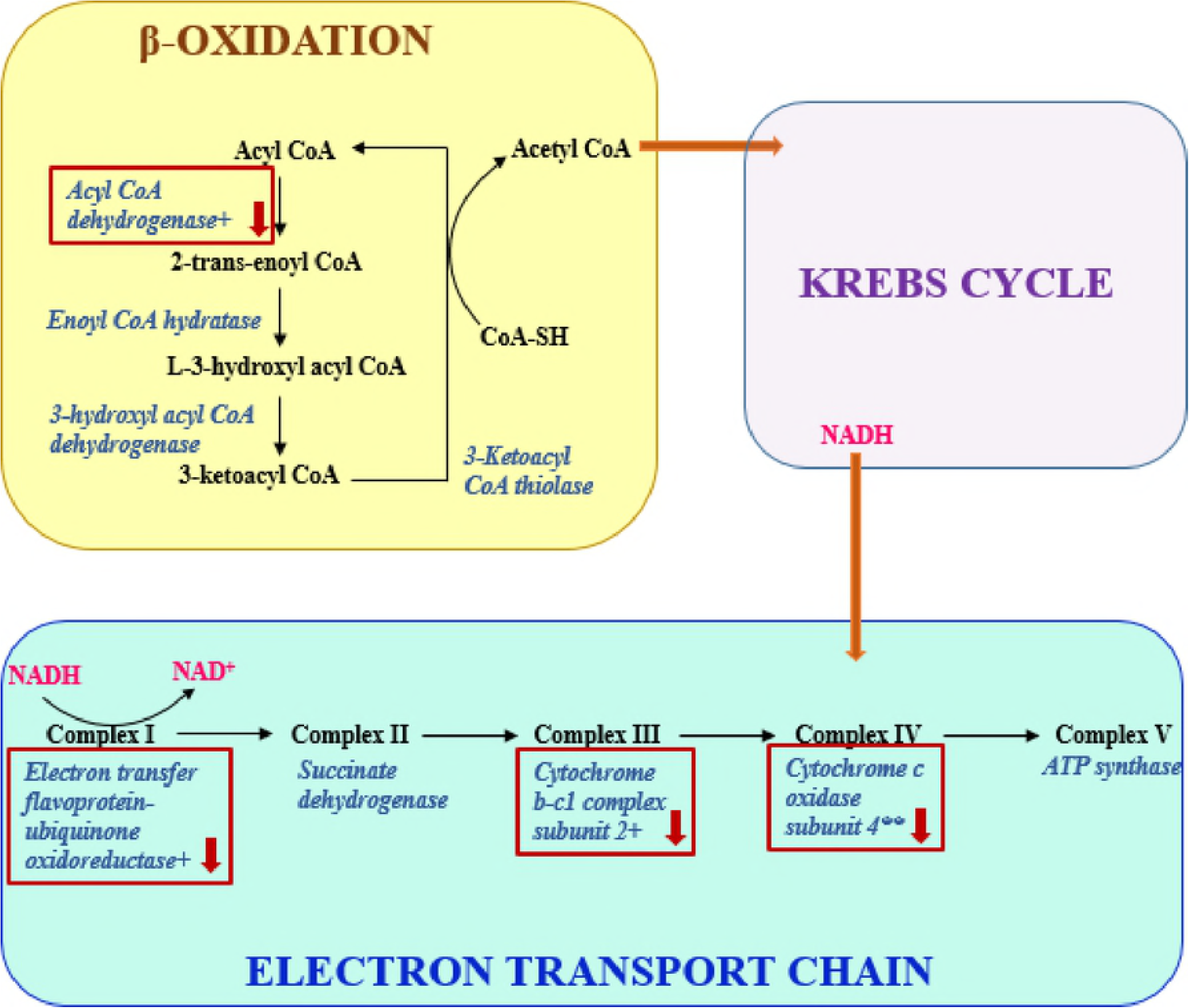
Graph of upregulated proteins in severe apnea group when compared to control. Values are measured as fold change.

Creatine kinase (CK) is found in high concentrations in tissues that consume ATP rapidly, such as the myocardium. Creatine kinase creates an “energy buffer” by phosphorylating via ATP, resulting in phosphocreatine and ADP. This reaction is reversible and in times of high energy demand phosphocreatine can convert ADP to ATP [17]. Clinically, CK has been associated with hypertension and myocardial ischemia and is used as a biomarker for myocardial infarction [18, 19]. This protein has also been shown to have increased serum concentrations in OSA sufferers who have no prior evidence of cardiovascular disease [20]. Many congestive heart failure studies have shown decreased levels of CK in both atria and ventricles, indicating decreased energy buffering effect [21, 22]. However, CK has been shown to remain at normal levels in models of compensated hypertrophy [19–21].

Aldose reductase (AR) is an important component of the polyol pathway and is responsible for converting glucose to sorbitol while oxidizing NADPH. The polyol pathway is an accessory pathway often activated when large concentrations of glucose are present in the cell and the main glycolytic pathway involving hexokinase becomes saturated, often due to hyperglycemia [23]. Increased dependence on the polyol pathway results in ATP scarcity and decreases the contractile efficacy of sarcoplasmic reticulum [24]. Experiments involving global ischemia and reperfusion illustrate how high concentrations of AR potentiate infarction and increases CK concentration in the myocardium [25]. Age has proven to be an important factor in determining infarct area in high AR models with global ischemia, but damage is significant at any age [25, 26].

Long chain acetyl CoA dehydrogenase (LACA) is essential for lipid metabolism and facilitates lipid integration into the citric acid cycle [27]. Cardiac hypertrophy and ischemia have been shown to increase glycolysis while decreasing lipid metabolism, resulting in metabolic characteristics of the fetal heart [28, 29]. Although chronic hypoxia is a significant problem for OSA sufferers, the cause of cardiomyopathy in OSA patients is often hypertension [2, 30]. Hypertension and hypertensive cardiomyopathy have both been related to a decrease in fatty acid oxidation and LACA levels [28, 31].

Heat shock protein 72 (HSP72), and other HSPs ensure normal protein folding and unfolding in periods of high stress, such as increased temperatures and ischemic conditions [32, 33]. Proper unfolding of denatured proteins that cannot be handled by chaperones often require degradation via HSPs like ubiquitin [33]. HSP72 has been associated with increased myocardial contractility when induced prior to cardiac stress and is shown to have a cardio protective effect in ischemia-reperfusion experiments [32, 34]. In the atria, HSP72 has proven effective in reversing atrial fibrillation in mitral valve surgery patients and reversing fibrosis of the atria induced by angiotensin II [35, 36]. Interestingly, HSP70 levels have been shown to decrease during sleep in OSA patients, whereas another study found an increase in daytime HSP70 levels when compared to control [37, 38].

Alpha-myosin heavy chain (MHCα) is a motor protein and one of the major driving forces in cardiac contraction. MHCα is the faster and more powerful MHC isoform, with MHCβ displaying a slower, energy conserving response [39, 40]. Atrial MHC in humans consists of 75% MHCα, and ventricles have 5% MHCα and 95% MHCβ [39]. Changes have been observed in MHC ratios in heart failure, with MHCα being downregulated in murine models and humans receiving heart transplants [41–43]. The downregulation of MHCα in heart failure may be slowed by exercise [41]. Typically, mammals with a faster resting heart rate and lower body mass possess higher levels of MHCα, and male humans have higher MHCα levels than females [39, 40].

Downregulated proteins in severe apnea samples when compared to control samples are shown in **Fig 4** and include cytochrome c oxidase subunit 4 isoform 1, mitochondrial precursor (-9.6 fold), peroxiredoxin 3 (-6.9 fold); troponin I, cardiac muscle (-3.3 fold), triosephosphate isomerase (-3.4 fold), ADP/ATP translocase 1 (-3.4 fold); Ba1-647 (-3.4 fold), L-lactate dehydrogenase B chain (-3.4 fold), malate dehydrogenase, cytoplasmic (-3.4 fold), and long-chain specific acyl-CoA dehydrogenase, mitochondrial precursor (-3.0 fold).

**Fig 4.**
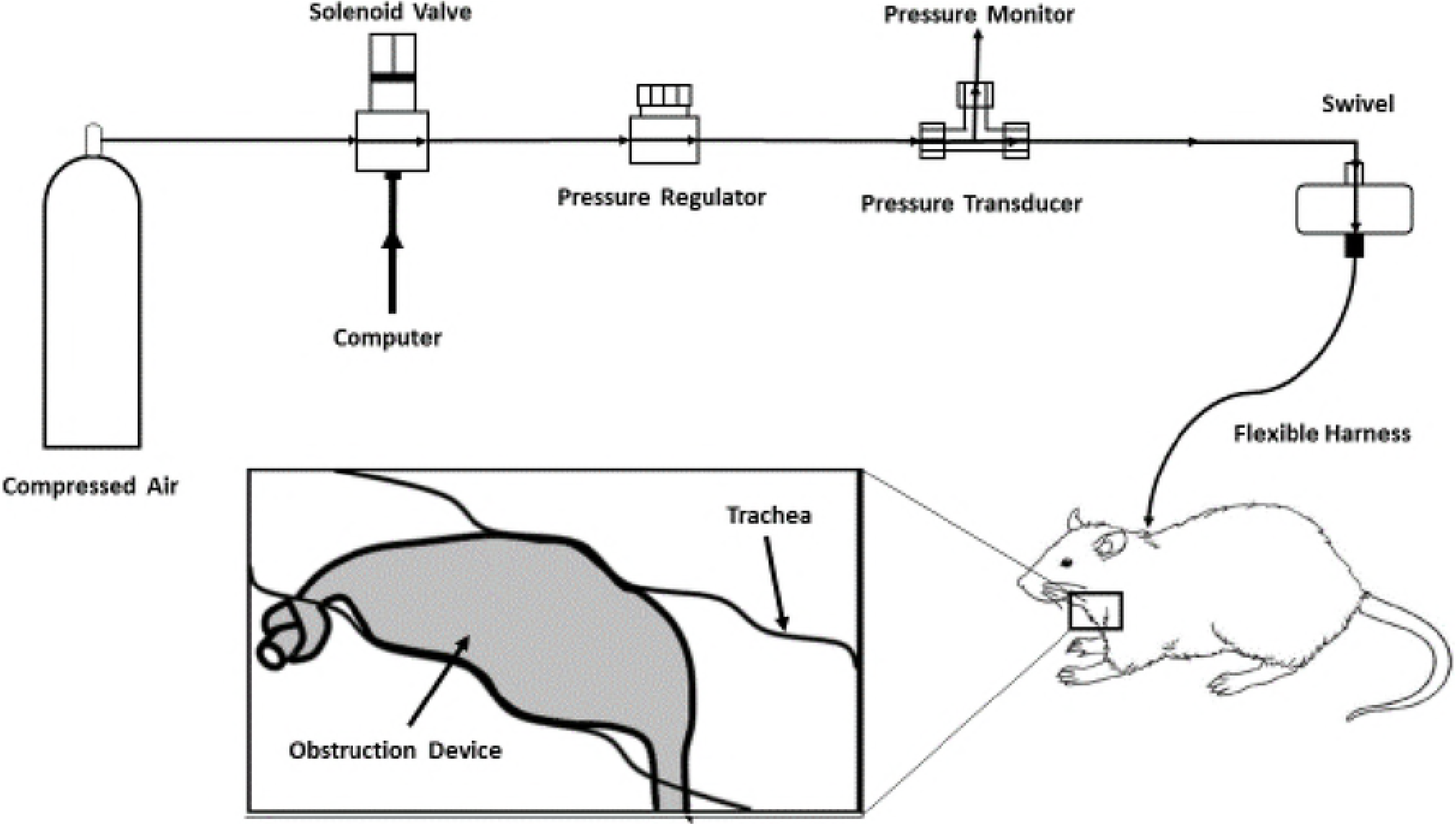
Graph of downregulated proteins in severe apnea group when compared to control. Values are measured as fold change.

Cytochrome c oxidase (CCO) is a large mitochondrial transmembrane protein located in the inner mitochondrial membrane and represents the final step in the electron transport chain [44]. The enzyme translocates electrons across the membrane to be combined with molecular oxygen to create water. During this process, CCO transfers protons across the mitochondrial membrane, into the intermembrane space, establishing an electromechanical potential for the creation of ATP [44, 45]. CCO is downregulated in models of heart failure and has been related to cardiac apoptosis in post-ischemic models [46, 47]. Oxidative stress has been shown to inhibit CCO independently of available electromechanical membrane potential and is also dependent on ATP levels as a feedback mechanism to decrease production of reactive oxygen species [48].

Peroxiredoxin 3 (P3) is a mitochondrial antioxidant that targets peroxides and is essential to redox signaling [49]. Increased peroxide levels have been observed in models of chronic intermittent hypoxia and ischemia-reperfusion, with increased oxidation of P3 [50, 51]. Similar increases in ROS have been observed in adults with OSA, resulting in upregulation of thioredoxin [52]. Peroxiredoxin 2 downregulation has been associated with atherosclerosis and upregulation of peroxiredoxin 2 is associated with a cardioprotective effect in ischemic models [53, 54].

ATP/ADP translocase (AAC) is a mitochondrial transmembrane protein responsible for the transportation of ATP and ADP to and from the cytosol for use and phosphorylation, respectively [55]. Decreases in AAC regulation correlate with decreases in ATP production and increases in mitochondrial permeability in ischemic and heart failure cardiomyocytes [56, 57]. Increased regulation of AAC has been shown to prevent apoptosis via the tumor necrosis factor β pathway, and increased peroxidative stress is related to decreased AAC regulation [57–59].

Citrate synthase plays an important role as the first enzyme in the citric acid cycle by converting oxaloacetate and acetyl CoA to citrate and CoA-SH [60]. As described previously, the citric acid cycle and mitochondrial metabolism as a whole is downregulated in various types of heart failure [28, 61, 62]. In addition, mitochondrial dysfunction and citrate synthase downregulation have been shown to be exacerbated by atrial fibrillation [63, 64].

Troponin T is an essential excitation-contraction coupling protein and is often used in the clinical setting as a biomarker for myocardial infarction [65, 66]. Increased serum concentrations of troponin T have also been linked to atrial fibrillation (AF) and tachycardia, and troponin increases in recently diagnosed AF patients predicts coronary artery disease [67–69]. Troponin T levels in serum has also been correlated with cardiomyopathy and increased OSA severity [70–72].

### Dysregulated proteins in moderate apnea samples, compared to controls

Upregulated proteins in moderate apnea samples when compared to controls are shown in **Fig 5**. Some of these proteins include Annexin V (6.3 fold), sarcalumenin, isoform CRA-a (5.1 fold), ATP synthase, mitochondrial F1 complex, alpha subunit (6.3 fold); ATP synthase beta subunit (4.4 fold), cytochrome b-c1 complex subunit 1, mitochondrial precursor (3.8 fold), tubulin beta-4B chain (3.3 fold), pyruvate dehydrogenase E1 component subunit beta, mitochondrial precursor (3.1 fold) o raspartate aminotransferase, mitochondrial (2.3 fold).

**Fig 5.**
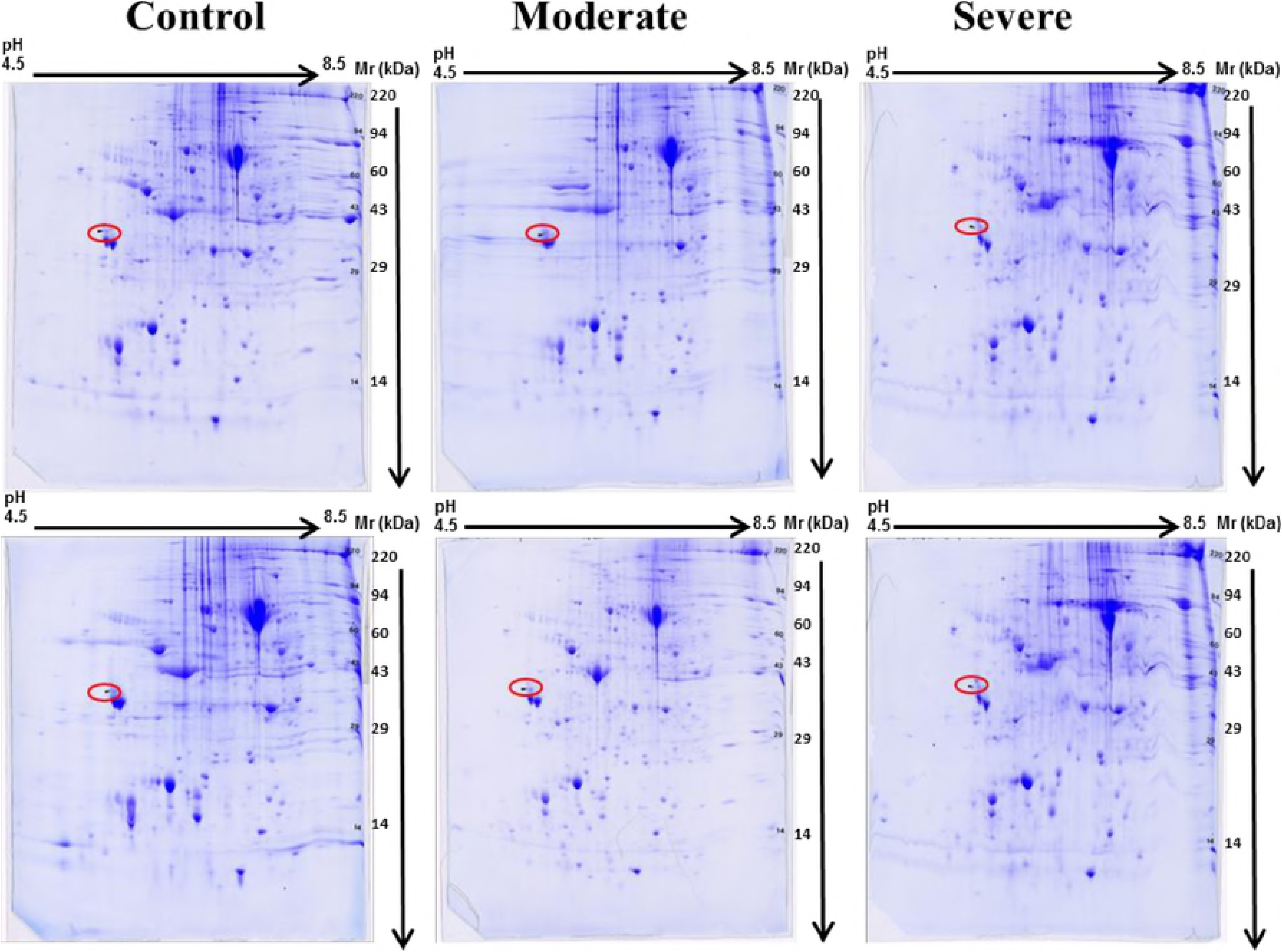
Graph of upregulated proteins in moderate apnea group when compared to control. Values are measured as fold change.

Annexin V is commonly used for detection of apoptotic processes using flow cytometry because of an affinity to phosphatidylserine [73]. Increased serum levels of annexin V have been observed in ischemia/reperfusion models, ventricular hypertrophy, and end stage cardiomyopathy patients [74, 75]. Annexin V has been associated with endothelial dysfunction and is a predictor of cardiac events [76]. Endothelial dysfunction and increased serum levels of annexin V have been observed in OSA patients, and is exacerbated during sleep [77–79]. Annexin V is associated with increased prevalence of atrial natriuretic peptide granules and has been shown to concentrate around the Z-line in rat atria [80, 81]. However, annexin V is decreased in the human atria suffering from atrial fibrillation [82].

Sarcalumenin (SAR) is located in the sarcoplasmic reticulum and is necessary for calcium buffering [83, 84]. SAR is located only in the atria and has been shown to be essential to cardiac function during aerobic and anaerobic metabolism; SAR knockout mice have exhibited decreased cardiac function [83, 85]. Hypertensive cardiomyopathy increases instances of spontaneous calcium release, and therefore displays predisposition to arrhythmia [86].

ATP synthase is an essential mitochondrial enzyme responsible for the phosphorylation of ADP by utilizing an electromechanical H^+^ gradient as an energy source. This molecular motor is responsible for the majority of ATP production in the cell [87]. Atrial fibrillation has been related to upregulation of noncoding RNA (AK055347), which directly correlates with ATP synthase regulation [88, 89]. Interestingly, isoflourane has been shown to have a cardioprotective effect in human atria, preconditioning mitochondria to perform with greater efficiency, benefiting the cell in ischemic conditions [90]. Similar effects have been observed with traditional ischemic preconditioning [91].

Tubulin β is a microtubule component of the cytoskeleton and upregulation is shown to disrupt cardiac β adrenergic stimulation, resulting in alteration of sodium current, leading to atrial arrhythmias such as sick sinus syndrome and life threatening tachyarrhythmia [92, 93]. Tubulin β is also upregulated in cardiac hypertrophy and impairs cardiac contractility [94, 95]. Regulation of mitochondrial respiration is closely related to tubulin β localization and regulation, mitochondrial creatine kinase is also suggested to interact with tubulin β [96–98].

cAMP-dependent protein kinase (PKA) is a component in G protein-coupled receptor (GPCR) signaling, specifically β adrenergic stimulation in the heart [99]. PKA has also been shown to interact with calcium signaling and is related to calcium induced pro-arrhythmic activity [100, 101]. Increased β adrenergic stimulation is often present in heart failure patients, with PKA levels and calcium activity also increased. This has made PKA an important pharmacological target for potential heart failure medications [102].

Downregulated proteins in moderate apnea samples when compared to controls are shown in **Fig 6** and include HSP 90-beta (-12.5 fold); long-chain specific acyl-CoA dehydrogenase, mitochondrial precursor (-6.9 fold), transcriptional adapter 2-alpha (-6.9 fold); alpha actinin 2 (-7.6 fold), cytochrome b-c1 complex subunit 2, mitochondrial precursor (-5.5 fold), vinculin, isoform CRA_a (-3.7 fold), NDRG1 related protein NDRG2a2 (-3.6 fold) and myristoylated alanine-rich C-kinase substrate (-4.0 fold).

**Fig 6.**
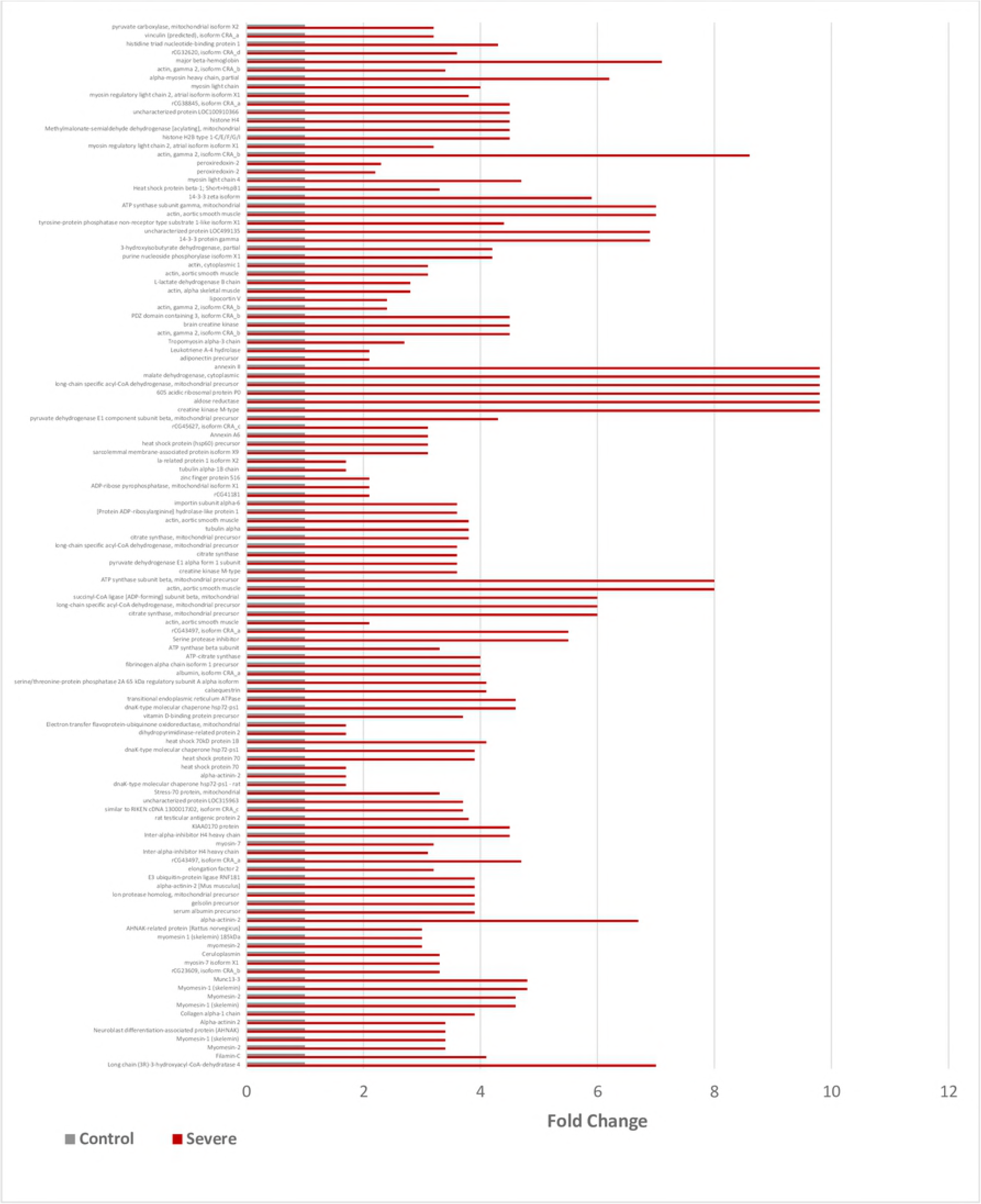
Graph of downregulated proteins in moderate apnea group when compared to control. Values are measured as fold change.

Vinculin is a focal adhesion protein responsible for biding α actin to integrin, acting as a junction and aiding cell rigidity and motility [103, 104]. Aging of the myocardium results in fibrosis and decreased cardiac contractility, with a correlating increase of vinculin [105]. Human mutations in vinculin and vinculin knockout mice have been related to increased incidences of dilated cardiomyopathy and sudden cardiac death [106, 107]. Vinculin absence has been shown to both facilitate contractile dysfunction and increase prevalence of ectopic beats in the myocardium, including ventricular tachycardia [106, 107].

NDRG1 is involved in stress response pathways by inhibiting cell growth and encouraging DNA repair, and is considered an important preventer of carcinogenesis [108, 109]. NDRG1 had been shown to be upregulated in hypoxic conditions and is increased in the p53 apoptotic pathway [110–112].

Myristoylated alanine-rich C-kinase (MARCKS) is a protein kinase C substrate that participates in actin crosslinking and interacts with calmodulin, a protein that participates in inflammatory and apoptotic mechanisms [113–115]. Protein kinase C and MARCKS have been shown to play important roles in cardiac cell differentiation and development in fetal and neonatal rats [116, 117].

Additionally, many enzymes associated with aerobic metabolic pathways like glycolysis, kreb’s cycle and anaerobic respiration were dysregulated in severe and moderate apnea when compared to control which was also identified to be dysregulated in our 1-D analysis [5].

### Dysregulated proteins in severe apnea compared to moderate apnea

Upregulated proteins in severe apnea samples when compared to moderate apneas are shown in **Fig 7** and include gelsolin precursor (10.6 fold), lon protease homolog, mitochondrial precursor (10.6 fold), E3 ubiquitin-protein ligase RNF181 (10.6 fold), Myomesin-1 (skelemin) (4.8 fold) and Myomesin-2 (4.8 fold).

**Fig 7.**
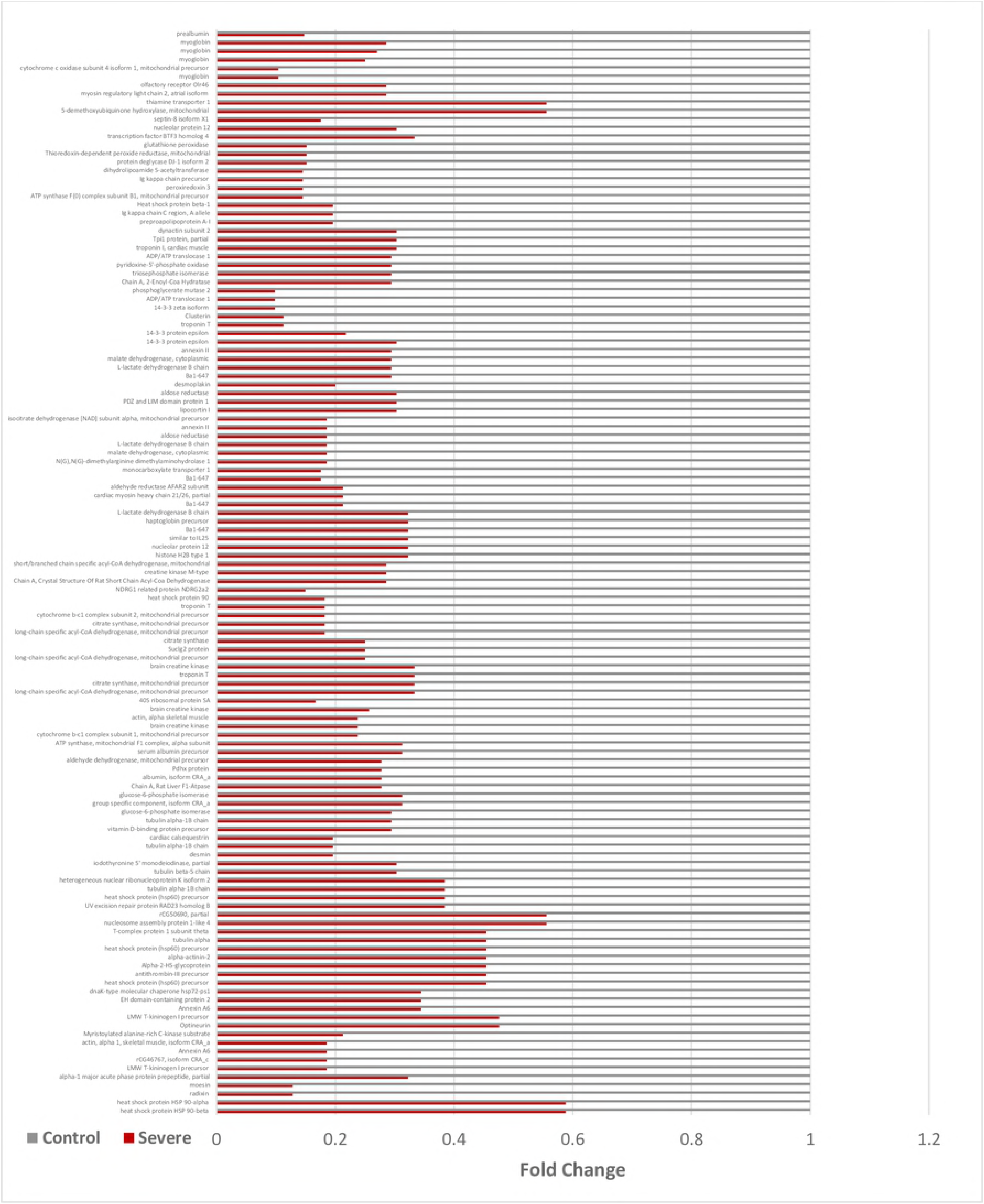
Graph of upregulated proteins in severe apnea group when compared to moderate apnea group. Values are measured as fold change.

Gelsolin is a protein responsible for cytoskeletal actin regulation and is Ca^2+^ dependent [118]. Expression of gelsolin is increased after myocardial injury, in mouse models of dilated and pressure-overload cardiomyopathy, and in end stage heart failure [118–120]. The increased expression of gelsolin has been related to Ca^2+^ disturbances in the diseased cardiomyocyte, whereas decreased atrial gelsolin concentration is related to increased atrial fibrillation prevalence in mice. [118, 121, 122].

Lon protease homolog (LONP1) is a mitochondrial protein responsible for the degradation of damaged mitochondrial proteins [123]. Increased levels of reactive oxygen species and hypoxia have been associated with mitochondrial adaptation via increased levels of LONP1, whereas decreased LONP1 levels have been associated with higher incidence of cell necrosis [123–125]. Oxidation of LONP1 in a mouse heart failure model resulted in decreased cardiac contractility and overall LONP1 dysfunction [126].

Ubiquitin protein ligases catalyze the transfer of ubiquitin from ubiquitin conjugating enzyme to a protein, marking the protein for transport, modification, or destruction via the proteasome [127]. Ubiquitin protein ligases are essential to the turnover of sarcomere proteins and certain ubiquitin ligases are responsible for myocardial atrophy, mitigating hypertrophy, exacerbating heart failure, protection against apoptosis, and mediating apoptosis [128, 129]. Due to the numerous varieties and complex roles of ubiquitin ligases in the myocardium, little conclusion can be drawn from differences in ubiquitin ligase regulation.

No significant downregulation was observed in severe apnea samples when compared to moderate apnea samples.

### Protein charge heterogeneity on 2D PAGE

Proteins expressed from a single gene can undergo different post-translational modifications (PTMs) on the amino acid side chains or at the C- or N- termini to give many protein products [130]. The PTMs include; acetylation, formylation, methylation, phosphorylation, nitrosylation and others. These modifications alter the pI of the proteins with a slight change in the molecular weight which will induce a spot shift in the 2D gels across the pH gradient [131–134]. Hence, the protein products can migrate to multiple spots on the gel. This can be observed in our study where we have found the same protein to be both upregulated and downregulated in severe and/or moderate when compared to control. For example, protein aldose reductase was found to be upregulated in severe when compared to control in spot no. 1516 (**S1 Table**) as well as downregulated in spot no. 1485 and 1489 (**S1 Table**). These different spots are observed for the same protein due to its charge heterogeneity caused by the PTMs. To identify the PTMs that are responsible for the differences between these two protein isoforms, we re-analyzed the data using more PTMs as variable modifications. However, the protein coverage was not sufficient to draw any specific conclusion at this time.

### Classification of Proteins

Dysregulated proteins were classified according to molecular function and biological processes using the PANTHER database and the results are depicted in **S3 and S4 Figs.** Severe and moderate apnea groups had catalytic enzymes dysregulated (45.3% and 44.4%, respectively) as well as binding proteins (35.9% and 38.9%, respectively). This can be seen from the list of proteins where several catalytic enzymes involved in aerobic respiration and also structural proteins involved in muscle contraction are dysregulated. Most dysregulated proteins participate in metabolic processes (severe: 24.1%, moderate: 29%), cellular processes (severe: 27.1%, moderate: 24.6%) and biogenesis (severe: 10.5%, moderate: 15.5%).

## Conclusion

The goal of the current study was to better characterize molecular changes that occur in the atria when subjected to hypoxia and decreased intrathoracic pressure, as in OSA. Apnea has also been found to cause myocardial dysfunction [135]. We used 2D-PAGE and LC-MS/MS analysis to further define proteomic dysregulation as it occurs in this model. We have discovered a number of mitochondrial proteins to be downregulated in severe and moderate apnea samples when compared to controls, indicating possible mitochondrial dysfunction. This correlates with studies on heart failure energetics, also indicating decreased mitochondrial efficacy and mitochondrial proteomic dysregulation [28, 64]. Increased regulation of structural proteins has been observed, such as tubulin and alpha myosin, as well as proteins that are involved in apoptotic mechanisms. Proteins involved in lipid metabolism, such as Long chain acetyl CoA dehydrogenase, have increased, contrary to current literature on fatty acid metabolism in the diseased heart [27–29]. Increased regulation of heat shock proteins were observed, indicating possible compensatory activity, although antioxidants such as peroxiredoxin 3 have been downregulated. Multiple proteins known for pro-arrhythmic interactions appear dysregulated, including structural, GPCR, and sarcomeric proteins [6, 69, 92, 100, 104]. Although many of the proteomic changes observed are indicative of cardiomyopathy and may enhance arrhythmias, this model does not match the hypoxic cardiomyopathy phenotype entirely, possibly due to additional factors related to airway obstruction and global hypoxia [21, 26, 28, 31]. Further research involving a larger number of samples per group may yield more consistent results and allow for the discovery of additional dysregulated proteins.

While in our previous study that employed 1D-PAGE followed by nanoLC-MS/MS analysis, we focused on a comparison of the full proteomes of the atrial tissues from control, moderate apnea and severe apnea,, the in the current study that employed 2D-PAGE followed by nanoLC-MS/MS analysis, we focused only on the proteins that are dysregulated in the atrial tissues from control, moderate apnea and severe apnea. Therefore, the two studies could not only confirm, but also complement each other and produced also a more accurate representation of the proteomic changes as they occur in the atria. This is also evidenced by the large number of dysregulated identified proteins in 2D-PAGE, which greatly complements our previous study [5]. Specifically, in the 1D-PAGE study we found that some of the enzymes involved in the aerobic (i.e. fructose-bis-phosphate aldolase, glyceraldehyde 3-phosphate dehydrogenase and phosphoglycerate kinase) and anaerobic (lactate dehydrogenase) glycolysis, as well as in the Krebs cycle (malate dehydrogenase and isocitrate dehydrogenase) are downregulated.

In the current 2D-PAGE study, additional enzymes involved in the aerobic glycolysis (i.e. phosphoglycerate mutase), Krebs cycle (succinylCoA ligase), electron transport chain (electron transport flavoprotein, a subunit of the Complex I and subunit 2 of the Complex III/cytochrome bc1 complex), anaplerotic reactions (enzymes involved in the replenishment of the metabolites (i.e. oxaloacetate, alpha-keto-glutarate, succinylCoA) involved in the Krebs cycle (aspartate aminotransferase and pyruvate carboxylase), and beta-oxidation (acyl-CoA dehydrogenase) were also found, suggesting that the OSA-induced oxygen deprivation of the heart induced downregulation of the entire glycolytic pathway, Krebs cycle, anaplerotic reactions, electron transport chain and beta-oxidation of the fatty acids (**Figs 8 and 9**).

**Fig 8:**
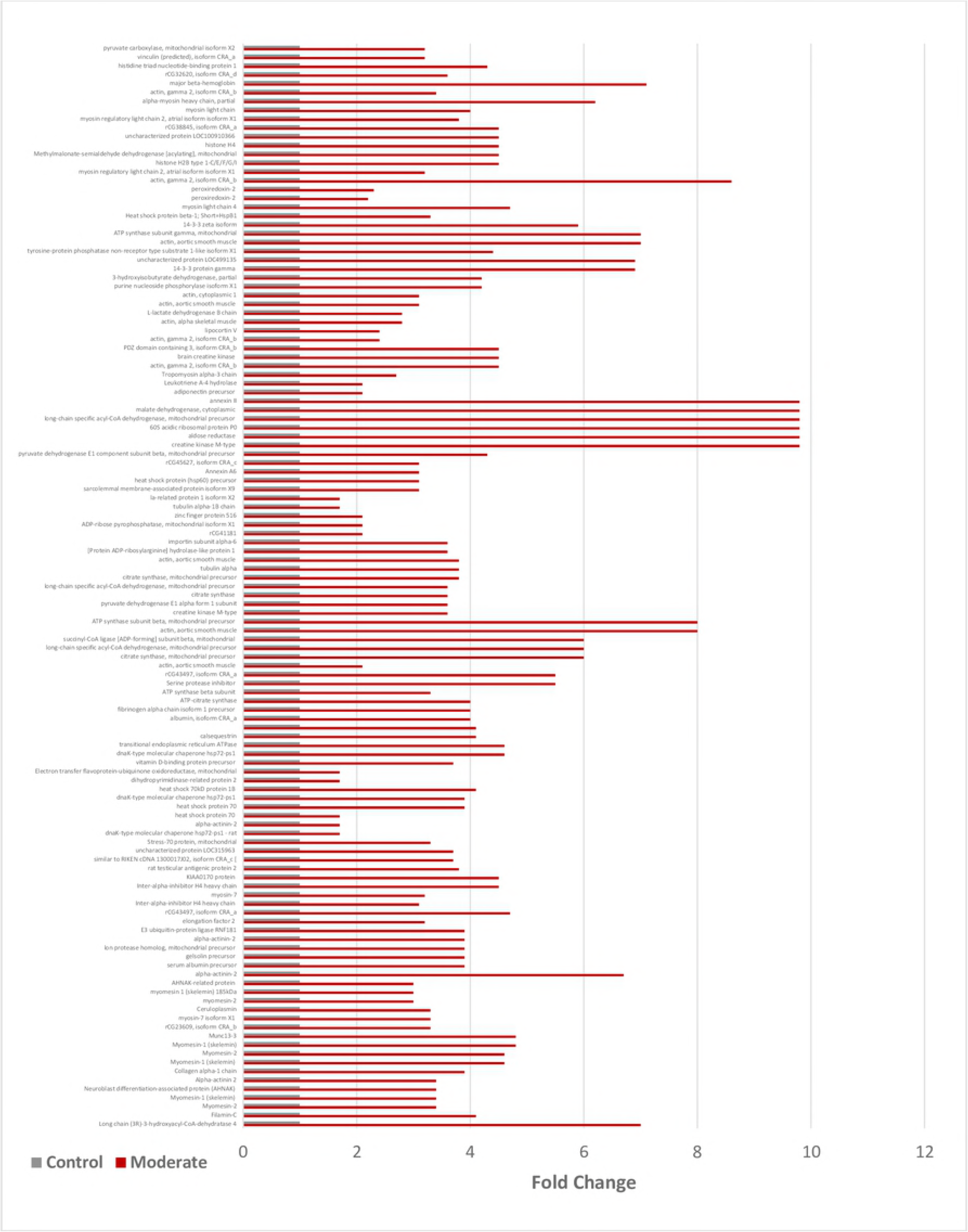
Outline of the aerobic (Glycolysis and Krebs cycle) and anaerobic (lactic acid fermentation) respiratory pathways depicting the downregulation of enzymes involved (shown in red downward arrow) in severe and moderate apnea when compared to control. The downregulated enzymes with an asterisk represents the findings from the 1-D proteomic results, enzymes with two asterisks represent the findings from the current 2-D proteomic results and the enzymes with a plus sign are found to be downregulated in both 1-D and 2-D proteomic analysis.

**Fig 9:**
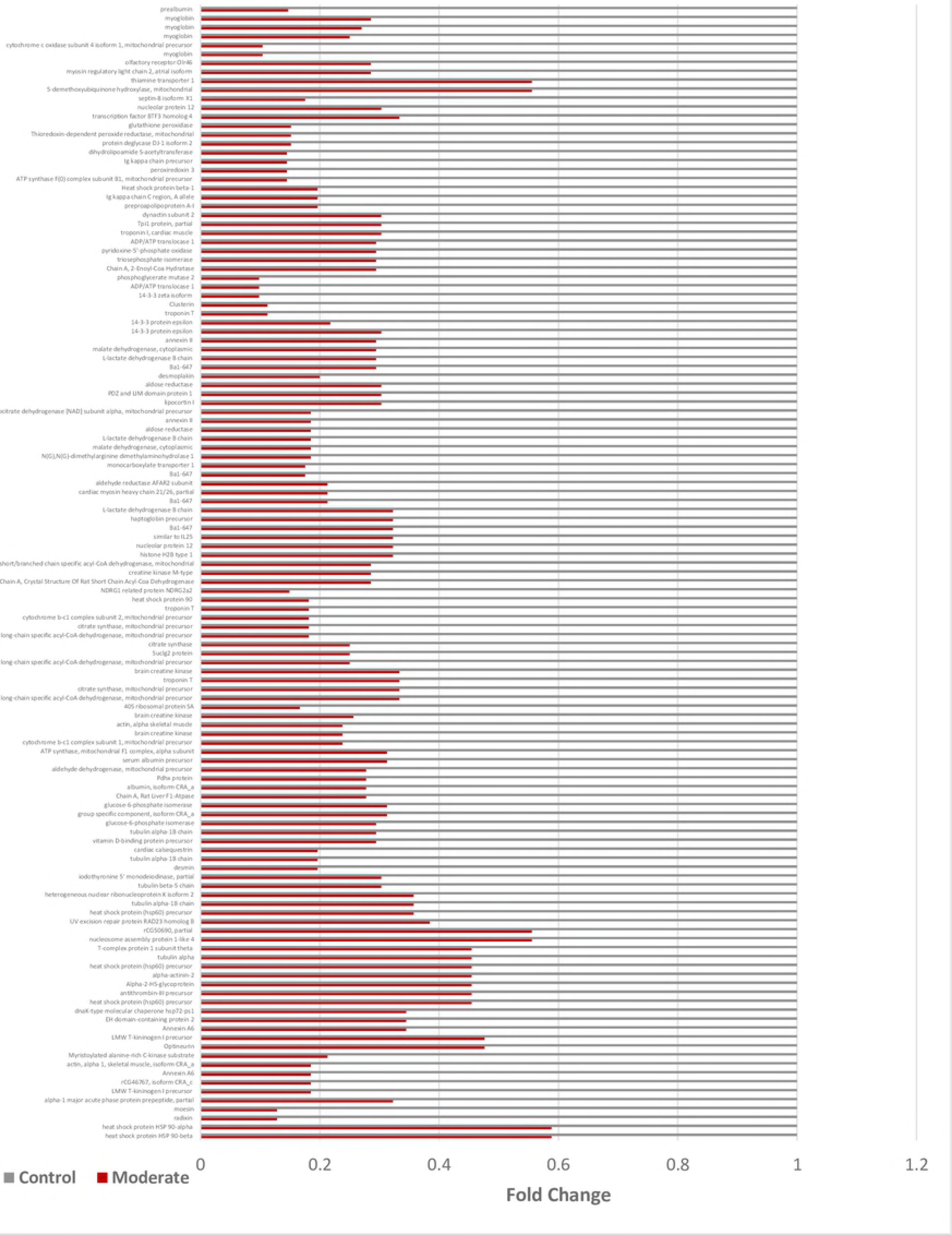
Depiction of the β-oxidation pathway and electron transport chain showing all the enzymes downregulated (shown in red downward arrow) in severe and moderate apnea condition when compared to control. The downregulated enzymes with an asterisk represents the findings from the 1-D proteomic results, enzymes with two asterisks represent the findings from the current 2-D proteomic results and the enzymes with a plus sign are found to be downregulated in both 1-D and 2-D proteomic analysis.

Overall, we believe our model accurately recreates the cardiac effects of OSA as it occurs in humans. This model also led to identification of protein dysregulations that may indicate a potentiation for atrial arrhythmia, decreased atrial compliance, increased inflammation, and risk of cardiomyopathy.

## Acknowledgements

The Authors thank Dr. Babu Suryadevara and the Clarkson’s Center for Advanced Materials Processing (CAMP) from Clarkson University and from the Masonic Medical Research Laboratory for supporting the initial work on this project through a Clarkson University-Masonic Medical Research Lab collaborative program. We would also like to thanks Kendrick Laboratories, Inc. for the 2D analysis which included computerized comparison of the gels, statistical analysis of the spots identified after 2D-PAGE.

The authors declare no conflicts of interest related to this work.

## Supporting Information

**S1 Fig. Images of control, moderate, and severe apnea silver stained 2D polyacrylamide gels.** The circles on the left side of each 2D polyacrylamide gel indicate the location of the IEF internal standard, tropomyosin with a Mr of 33,000 and pI of 5.2.

**S2 Fig. Depiction of averaged severe apnea samples compared to control samples.** Blue circled spots represent increased polypeptide regulation in severe apnea groups, red circled spots indicate peptide downregulation in severe apnea versus control.

**S3 Fig. Classification of dysregulated proteins in severe OSA samples organized by molecular function and biological process as charicterized by PANTHER**.

**S4 Fig. Classification of dysregulated proteins in moderate OSA samples organized by molecular function and biological process as characterized by PANTHER**.

**S1 Table. List of proteins identified from the 2D-PAGE gel spots which were dysregulated in severe apnea when compared to control.** A positive fold change indicates the upregulation of protein and a negative fold change indicates the downregulation of protein in severe apnea when compared to control.

**S2 Table. List of proteins identified from the 2D-PAGE gel spots which were dysregulated in moderate apnea when compared to control.** A positive fold change indicates the upregulation of protein and a negative fold change indicates the downregulation of protein in moderate apnea when compared to control.

**S3 Table. List of proteins identified from the 2D-PAGE gel spots which were dysregulated in severe apnea when compared to moderate.** A positive fold change indicates the upregulation of a protein.

